# Mutational pressure drives differential genome conservation in two bacterial endosymbionts of sap feeding insects

**DOI:** 10.1101/2020.07.29.225037

**Authors:** Gus Waneka, Yumary M. Vasquez, Gordon M. Bennett, Daniel B. Sloan

## Abstract

Compared to free-living bacteria, endosymbionts of sap-feeding insects have tiny and rapidly evolving genomes. Increased genetic drift, high mutation rates, and relaxed selection associated with host control of key cellular functions all likely contribute to genome decay. Phylogenetic comparisons have revealed massive variation in endosymbiont evolutionary rate, but such methods make it difficult to partition the effects of mutation vs. selection. For example, the ancestor of auchenorrhynchan insects contained two obligate endosymbionts, *Sulcia* and a betaproteobacterium (*BetaSymb*; called *Nasuia* in leafhoppers) that exhibit divergent rates of sequence evolution and different propensities for loss and replacement in the ensuing ∼300 Ma. Here, we use the auchenorrhynchan leafhopper *Macrosteles sp. nr. severini*, which retains both of the ancestral endosymbionts, to test the hypothesis that differences in evolutionary rate are driven by differential mutagenesis. We used a high-fidelity technique known as duplex sequencing to measure and compare low-frequency variants in each endosymbiont. Our direct detection of *de novo* mutations reveals that the rapidly evolving endosymbiont (*Nasuia*) has a much higher frequency of single-nucleotide variants than the more stable endosymbiont (*Sulcia*) and a mutation spectrum that is even more AT-biased than implied by the 83.1% AT content of its genome. We show that indels are common in both endosymbionts but differ substantially in length and distribution around repetitive regions. Our results suggest that differences in long-term rates of sequence evolution in *Sulcia* vs. *BetaSymb*, and perhaps the contrasting degrees of stability of their relationships with the host, are driven by differences in mutagenesis.

**SIGNIFICANCE STATEMENT:** Two ancient endosymbionts in the same host lineage display stark differences in genome conservation over phylogenetic scales. We show the rapidly evolving endosymbiont has a higher frequency of mutations, as measured with duplex sequencing. Therefore, differential mutagenesis likely drives evolutionary rate variation in these endosymbionts.

## INTRODUCTION

Sap feeding insects in the order Hemiptera feed exclusively on nutrient deficient plant-saps (phloem and xylem) and rely on bacterial endosymbionts to provide lacking essential amino acids and vitamins (Buchner 1965; Sandström and Moran 1999). Such endosymbionts are housed intracellularly in specialized organs called bacteriomes and are subject to strict vertical transmission from mother to offspring. This repeated transmission bottlenecking and confinement within a host lineage results in genetic drift and endosymbiont genome decay (Moran 1996; Mira and Moran 2002; McCutcheon and Moran 2012; Bennett and Moran 2015; McCutcheon et al. 2019).

Ancient endosymbiont genomes are characterized by their extreme nucleotide composition bias and small size. In fact, many of these genomes are among the smallest known, lacking genes considered essential in free living bacteria (e.g., cell envelope biogenesis, gene expression regulation, and DNA replication/repair; McCutcheon and Moran 2012; Moran and Bennett 2014). Ancient endosymbiont genomes experience rapid sequence evolution and extensive lineage-specific gene deletions, leaving only a set of ‘core’ genes necessary to sustain a functioning nutritional symbiosis (Moran et al. 2009; Bennett, McCutcheon, et al. 2016). At the same time, these genomes exhibit remarkable structural stability, often displaying perfectly conserved synteny between lineages that diverged 10s to 100s of millions of years ago (McCutcheon et al. 2009a).

Missing DNA replication and repair genes, and conditions of relaxed selection, allow the AT mutation bias (which is common to most bacteria; Hershberg and Petrov 2010; Hildebrand et al. 2010; Long, Sung, et al. 2018) to drive AT content to levels above 75% in many endosymbionts (Moran et al. 2009; Moran and Bennett 2014; Wernegreen 2016). A notable exception is the GC rich genome of *Candidatus* Hodgkinia cicadicola (hereafter *Hodgkinia*; 58.4% GC), an endosymbiont of cicadas with an extremely small genome (144 kb) (McCutcheon et al. 2009b). However, population level estimates of the *Hodgkinia* mutation spectrum reveal an AT mutation bias, indicating that GC prevalence in *Hodgkinia* likely results from selection on nucleotide identity (Van Leuven and McCutcheon 2012).

In extreme cases of genome decay, ancient endosymbionts can go extinct when they are outcompeted and replaced by newly colonizing bacteria or yeast (Matsuura et al. 2018; McCutcheon et al. 2019). Comparisons amongst ancient endosymbionts reveal massive variability in genome stability and extinction rates. Insects in the suborder Auchenorrhyncha maintain two bacterial endosymbionts: “*Candidatus* Sulcia muelleri” (hereafter *Sulcia*) and a betaproteobacterium (hereafter *BetaSymb*), which are both thought to be ancestral to the group (∼300 Ma) (Moran et al. 2005; Dietrich 2009; Bennett and Moran 2013; Bennett and Mao 2018). The partner endosymbionts provision distinct and complementary sets of the 10 essential amino acids that animals are unable to synthesize and are unavailable in the host diet. *Sulcia* has been nearly universally retained over the ∼300 million year diversification of the Auchenorrhyncha (∼40,000 described species); in contrast, *BetaSymb* has been replaced in at least six major lineages, highlighting its relative instability (Dietrich 2009; Bennett and Moran 2015).

The apparent unreliability and relatively frequent loss of the *BetaSymb* lineage is mirrored by rapid rates of molecular evolution. Pairwise divergence estimates for the two ancestral symbionts in closely related leafhopper species (family *Cicadellidae*) reveal that *Sulcia* genomes are highly similar (99.68% nucleotide identity), while *BetaSymb* genomes are 30-fold more divergent (90.47% nucleotide identity) (Bennett, Abbà, et al. 2016). The dramatic differences in *Sulcia* and *BetaSymb* rates of molecular evolution have also been documented across more divergent auchenorrhynchan lineages (Bennett and Moran 2013; Mao et al. 2017; Bennett and Mao 2018).

While phylogenetic comparisons have greatly improved our understanding of the changes that endosymbiotic genomes experience, they do not adequately parse the relative contributions of mutagenesis, genetic drift, and natural selection. Although it is possible that selection differentially constrains sequence change in *Sulcia* and *BetaSymb* (Wernegreen 2016), it has been suggested that the low rate of evolution in *Sulcia* may correlate with decreased DNA replication (Silva and Santos-Garcia 2015) or low mutational input (Bennett et al. 2014). However, these hypotheses are difficult to test directly as traditional approaches for measuring bacterial mutation, such as mutation accumulation lines (Kucukyildirim et al. 2016; Long, Miller, et al. 2018), are not feasible for obligate endosymbionts. In one case, researchers were able to use the recent demographic history of the insect host to infer endosymbiont mutation rates (Moran et al. 2009), but the challenge of directly measuring mutation in endosymbionts continues to inhibit efforts to disentangle mutagenesis, drift, and selection in endosymbionts.

A direct measurement of mutation would ideally capture all variants, before some can be filtered by natural selection. While modern DNA sequencing technologies have facilitated an enormous increase in sequenced endosymbiont genomes, they have not necessarily translated to improvements in measuring *de novo* mutation in endosymbiont genomes. This is because typical sequencing error rates are too high (often above 10_-3_ errors per base pair [bp]) to detect rare variation in DNA samples (Schirmer et al. 2016). As a result, these methods can only detect single-nucleotide variants (SNVs) that have existed long enough to rise to a substantial frequency within the sample, either via drift or positive selection.

Fortunately, the recent advent of several high-fidelity DNA sequencing techniques provides the opportunity to obtain a snapshot of extremely low frequency SNVs in endosymbiont DNA (Sloan et al. 2018). One technique, called duplex sequencing, is particularly suited for such a measurement because it has an exceptionally low error rate of less than 2×10^−8^ errors per bp (Kennedy et al. 2014; Wu et al. 2020). The high accuracy of this method is achieved by tagging each original DNA fragment with random barcodes and producing multiple sequencing reads from both strands to obtain a consensus sequence. Importantly, this method is robust to single-stranded DNA damage and individual errors introduced during PCR amplification or sequencing because it separately tracks reads originating from the two complementary strands of the original biological molecule and requires support from each.

Here, we measured low frequency variants in the endosymbionts of the leafhopper (*Macrosteles*. sp. nr. *severini*) to test the hypothesis that differences in mutagenesis may be responsible for differential evolutionary stability between *Sulcia* and *BetaSymb* (called “*Candidatus* Nasuia deltocephalinicola” in leafhoppers; hereafter *Nasuia*). We assembled *Sulcia* and *Nasuia* reference genomes from a population of *M*. sp. nr. *severini* isolated from Hawaii, using standard Illumina and Oxford Nanopore (MinION) libraries. We then created duplex libraries, which were mapped to the reference genomes to detect *de novo* variants. We found a dramatically higher frequency of independent SNVs in *Nasuia* than in *Sulcia*, which suggests the elevated rate of evolution observed in *Nasuia* over phylogenetic scales is driven, at least in part, by large differences in mutagenesis

## MATERIALS AND METHODS

### Growth and DNA extractions of *Macrosteles sp. nr. severini* lines

A laboratory stock population of field collected *Macrosteles* sp. nr. *severini* from Sumida Farms, Aiea, Oahu Island, Hawaii was established on December 8, 2016. Identification of this species, which has not been formally described, followed Le Roux and Rubinoff (2009). For this experiment, eight laboratory populations were established on barley plants from a single randomly selected foundress on June 17, 2017. Populations were grown with occasional plant replacements to maintain colony health for approximately six months and harvested December 18-22, 2017. Two populations, referred to as A and B, survived to produce sufficient number of individuals for downstream genomic sequencing. Populations were further subdivided into two replicates each (A1, A2, B1, B2) for dissection and sequencing. For each replicate, bacteriomes from approximately 100 individuals were dissected out, pooled, and stored in 95% ethanol. Total DNA was extracted using a DNeasy Blood and Tissue kit (Qiagen) following manufacturer’s protocol with a 12-hour Proteinase K digestion.

To ensure variants detected with duplex sequencing were not an artifact of endosymbiont DNA transferred to the *M*. sp. nr. *severini* genome nuclear genome, we sequenced DNA isolated from *M*. sp. nr. *severini* heads, which should contain no true endosymbiont DNA. DNA was extracted from a pool of 20 dissected heads using the Qiagen Dneasy Blood and Tissue Kit.

### Construction of shotgun Illumina libraries

Standard shotgun Illumina libraries were created from the most concentrated bacteriome DNA sample (B1) as well as the head DNA sample using the NEBNext Ultra II FS DNA Library Prep Kit. We used 50 ng of input DNA, with 15-minute and 10-minute fragmentation steps for the bacteriome and head DNA, respectively. Samples were amplified with 4 cycles of PCR. The libraries had average lengths of 318 bp and 319 bp for the bacteriome and head DNA, respectively.

### Construction of duplex libraries from bacteriome DNA for variant detection

Separate duplex sequencing libraries were generated for each of the four replicate bacteriome DNA samples (A1, A2, B1, and B2). Duplex library preparation followed protocols described elsewhere (Wu et al. 2020). Briefly, samples were fragmented with the Covaris M220 Focused-Ultrasonicator and subsequently end-repaired (NEBNext End Repair Module), A-tailed (Klenow Fragment enzyme, 1 mM dATP), adapter ligated (NEBNext Quick Ligation Module) and treated with a cocktail of three repair enzymes (NEB CutSmart Buffer, Fpg, Uracil-DNA Glycosylase, Endonuclease III). 50 pg of repaired and adapter-ligated products were amplified for 19 cycles with the NEBNext Ultra II Q5 Master Mix (New England Biolabs M0544) and dual-indexed with custom IDT Ultramer primers. Adapter dimers were detected when DNA was assessed on an Agilent TapeStation 2200 (High Sensitivity D1000 reagents) and subsequently removed with size selection on a 2% BluePippin gel (Sage Science), using a target range of 300-700 bp. The pooled duplex libraries had an average length of 386 bp.

### Illumina sequencing of shotgun and duplex libraries

Shotgun and duplex libraries were sequenced on a NovaSeq 6000 platform (2×150bp) at the University of Colorado Cancer Center in separate sequencing runs. Sequencing resulted in 9.4 M read pairs for the bacteriome shotgun library. The shotgun library from *M*. sp. nr. *severini* head DNA was sequenced in two runs (the first run did not produce enough reads), which resulted in a total of 359 M read pairs. Duplex libraries were also sequenced in two runs (again, the first round of sequencing did not produce enough reads), which resulted in 37.7 M to 44.9 M read pairs per library. The raw sequencing reads are available via the NCBI Sequence Read Archive (SRA) under accessions SRR12112868 (A1), SRR12112867 (A2), SRR12112866 (B1), SRR12112865 (B2), SRX8635374 (*M*. sp. nr. *severini* head tissue) and SRR12112864 (B1: shotgun).

### MinION library construction and sequencing

To aid in genome assembly, we also sequenced the (B1) bacteriome sample with the Oxford Nanopore MinION (Jain et al. 2016), with 150 ng of input DNA on a FLO-MINSP6 flow cell, using the Rapid Sequencing Kit (SQK-RAD004). Data were processed using the MinION software release v19.10.1 with default MinKNOW parameters except that base calling was set to high accuracy mode. Only the first 1.1 M reads were used for genome assembly (which accounted for ∼ 2/3 of the data generated by the run and averaged 2972 bp in length). The resulting reads are deposited in the NCBI SRA under accession SRR12112863.

### Genome assembly and characterization

Endosymbiont genomes were assembled from the bacteriome Illumina shotgun and MinION libraries with the SPAdes genome assembler (v3.11.1) using the -nanopore flag (Bankevich et al. 2012). Scaffolds returned by SPAdes were searched (blastn v2.9.0+) against available *Nasuia* and *Sulcia* genomes from *M. quadrilineatus* (GenBank: CP006059.1 and CP006060.1, respectively) (Bennett and Moran 2013), and the two longest scaffolds were identified as near complete matches to *Sulcia* and *Nasuia*.

Evidence of chromosome circularity was assessed by searching for the scaffold beginning and ending sequences in the shotgun and MinION reads. Reads that contain the sequence of both scaffold ends (and therefore span the circular gap) were identified for both genomes. In both cases, the scaffolds ended in tandem repeat regions where the number of repeat units varied in different spanning reads (repeat length heterogeneity).

In the *Sulcia* assembly, the scaffold was broken at a 6-bp microsatellite (corresponding to positions 67046-67105 in the final assembly), with anywhere from eight to 11 repeat units in different spanning reads. The most common repeat number in spanning reads was 10, and the genome was manually adjusted accordingly. The same approach was used to confirm the most common tandem repeat number for three other regions in the *Sulcia* assembly that exhibited length heterogeneity (corresponding to regions 16069-16134, 174128-174181 and 178159-178299 in the final assembly), which were identified as scaffold breakpoints or ambiguous calls (Ns) in a separate SPAdes assembly run without the MinION data.

In the *Nasuia* genome, the repeat structure that broke the assembly is far more complicated, and its resolution was only possible through the use of the long-read MinION data (fig. 1). In spanning sequences, we found seven to 53 repeats of a ∼72 bp sequence (which contains only one G and one C), with 22 repeats being the most common repeat number. A complicating factor is the intermittent presence of an additional 7-bp sequence, which itself is a tandem repeat of the first seven bp of the 72-bp repeat, resulting in a mix of 72-bp and 79-bp repeats. The shotgun Illumina data were used to determine that 53.3% of repeats were 72 bp and 46.7 % of repeats were 79 bp. The spanning MinION read of median length, which contained 22 repeats total, was selected as the basis for resolving the *Nasuia* genome gap (fig. 1).

**Fig. 1.**
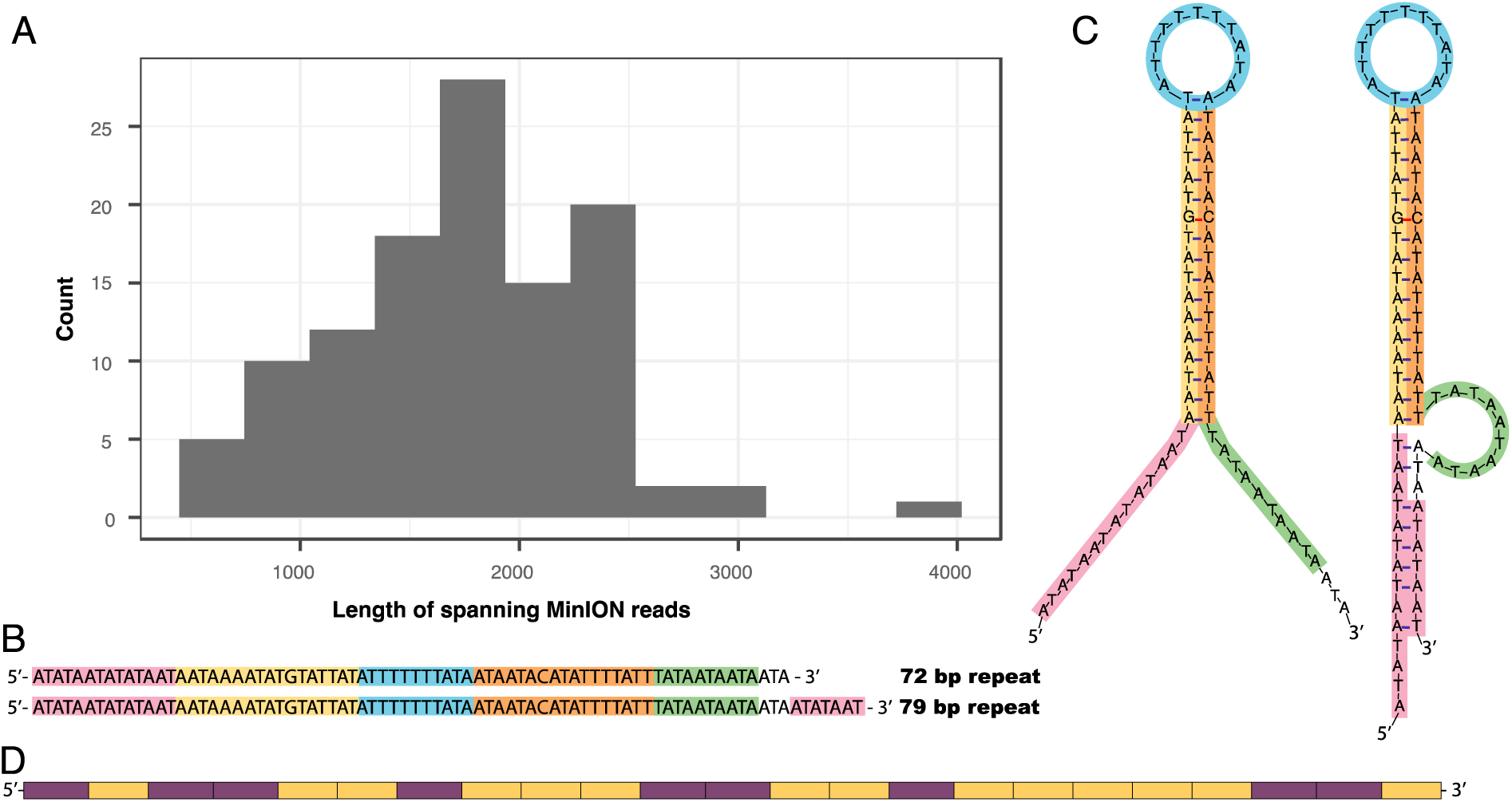
Characterization of a repetitive region in the *Nasuia-MSEV* genome. A repetitive region in the *Nasuia-MSEV* genome that could not be assembled with short read (150 bp) shotgun sequencing was reconstructed with long MinION reads. We detected massive repeat length variation in the 113 MinION reads that span the repeat region (i.e. reads that contain an element of the repeat and are flanked by unique identifier sequences on either end), as demonstrated in the histogram of read lengths **(A)** (min 500 bp, max 3775 bp, median 1757 bp). We determined that repeat units consist of either a 72 bp or 79 bp sequence **(B)** which are identical except for the 7 bp absent from the shorter sequence, and both contain an inverted repeat which could potentially form a stem loop as predicted by mfold (Zuker 2003) **C)**. In addition to the noted repeat number heterogeneity, we observed high variation in the relative distribution of 72 bp and 79 bp repeats. We chose the repeat order and repeat number based on the read with the median length. In **(D)** we present the number and order of 72-bp repeats (yellow) and 79-bp repeats (purple) in the read with the median length, and our final assembly.

**Fig. 2.**
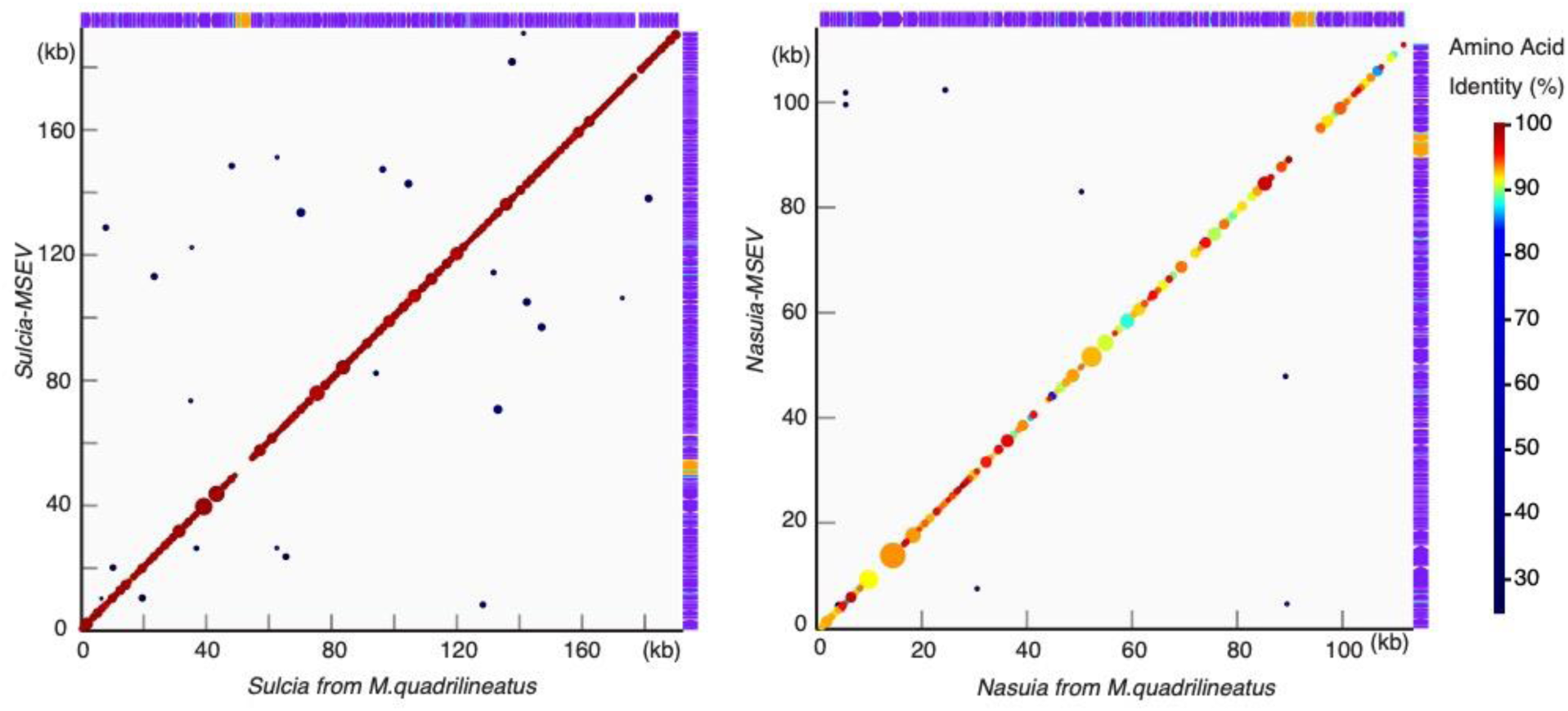
Synteny and sequence similarity of *Nasuia* and *Sulcia* from two closely related leafhopper species. Synteny plots reveal gene order to be perfectly conserved in both *Nasuia* and *Sulcia* of *M. quadrilineatus* and *M*. sp. nr. *severini*. Circles represent significant pairwise alignments between protein-coding genes in the two genomes, with circle diameter indicating alignment length. *Nasuia* exhibits a 700-fold higher sequence divergence than *Sulcia*, which is reflected in the lower amino acid identity (color scale on figure right) of *Nasuia* protein coding sequences. The compact nature of both genomes can be observed in the gene tracks on the top (*M. quadrilineatus* associated) and right (*M*. sp. nr. *severini* associated) of the synteny plots, where protein coding-genes are shown in purple, rRNA genes are in yellow, tRNA genes are in light blue, and intergenic regions are white. This figure was generated with GenomeMatcher (Ohtsubo et al. 2008)

After the most common repeat numbers were determined and circularity was confirmed, the breakpoints of the chromosomes were shifted, and the *Nasuia* scaffold was reverse complemented to reflect the orientation and genome positions of *Nasuia* and *Sulcia* accessions from the closely related *M. quadrilineatus* (Bennett and Moran 2013). We then aligned our new assemblies to *M. quadrilineatus* accessions using MAFFT (v7.453) under default parameters (Katoh and Standley 2013). Gaps in the alignments were tabulated with a custom script. Pairwise distances were calculated with MEGA (v10.1.8) (Kumar et al. 2018), using the Tamura 3-parameter model and a gamma distribution. Repetitive sequences were identified with custom scripts that report the position and length of homopolymers and microsatellites of 4+ bp in length (supplementary table 1) (Temnykh et al. 2001).

Both *Sulcia* and *Nasuia* were annotated with initial protein-coding gene predictions using Glimmer3 in Geneious v11.1.5 and were then checked against existing symbiont genomes from two other *Macrosteles* host species (Delcher et al. 2007; Bennett, Abbà, et al. 2016). All RNA annotations were done using the “annotate from reference” function in Geneious with a match threshold of >80% similarity. *Nasuia* and *Sulcia* genomes from *M. quadrilineatus* were used as the reference genome (Bennett and Moran 2013).

### Variant detection with duplex data

Duplex libraries were processed with our custom data analysis pipeline (Wu et al. 2020; https://github.com/dbsloan/duplexseq), which uses the random barcodes on read ends to pair raw reads into families with a single duplex consensus sequence (DCS). Consensus sequences were subsequently mapped against the newly assembled reference genomes and the resulting alignment files were parsed to identify indels, single nucleotide variants (SNVs), and coverage per bp. Supplementary table 2 reports total DCS coverage and percent mapping to endosymbiont genomes for the four replicates. A filtering step then checked for the presence of identified variants in a *k*-mer database (*k* = 39) created from the *M*. sp. nr. *severini* head DNA shotgun library (KMC v. 3.0.0; https://github.com/refresh-bio/KMC) to ensure variants are not derived from bacterial DNA that has been transferred to the host genome (Hotopp et al. 2007; Nikoh et al. 2010). All variants had counts well below those detected for *M*. sp. nr. *severini* nuclear and mitochondrial sequences.

## RESULTS

### Higher divergence in *Nasuia* than in *Sulcia* derived from closely related leafhoppers

We assembled *Nasuia* and *Sulcia* genomes (GenBank accession SAMN15504451 and SAMN15504450, respectively), using a combination of short-read (Illumina) and long-read (MinION) sequences from *M*. sp. nr. *severini* bacteriome DNA. The resulting genomes (referred to hereafter as *Nasuia-MSEV* and *Sulcia-MSEV*) were generally comparable to those previously sequenced from other hosts in terms of size (*Nasuia-MSEV*: 113 kb, *Sulcia-MSEV*: 190 kb), gene count (*Nasuia-MSEV*:173, *Sulcia-MSEV*: 224), and GC content (*Nasuia-MSEV*: 16.9%, *Sulcia-MSEV*: 24.0%).We aligned our new assemblies to the previously published *M. quadrilineatus* accessions and found pairwise distances of 7.68% and 0.01% for the *Nasuia* and *Sulcia* pairs, respectively (fig. 2). The ∼700-fold higher pairwise divergence in *Nasuia* than in *Sulcia* is much larger than 30-fold difference previously shown in a pairwise comparisons between endosymbionts of the closely related *M. quadrilineatus* and *M. quadripunctulatus* (Bennett, Abbà, et al. 2016). This disparity is driven by the near perfect sequence conservation in *M*. sp. nr. *severini* and *M. quadrilineatus Sulcia* genomes, which only differ at approximately one of every 10,000 positions. Transitions are responsible for 79.6% of the observed pairwise distance in *Nasuia*. The extremely small number of changes in the *Sulcia* comparison prohibits a meaningful calculation of the contribution of transitions to pairwise distance.

We found 1.00 and 0.28 gaps per kb in the *Nasuia* and *Sulcia* pairs, respectively (supplementary table 3). A large (1,352 bp) gap in the *Nasuia* pairs (supplementary table 3) corresponds to the mix of 72 and 79 bp repeats (22 total) in the *Nasuia-MSEV* assembly that we characterized with the MinION data (fig. 1). The lack of this region in the *Nasuia* accession from *M. quadrilineatus* likely reflects differences in methods of genome assembly, rather than an actual difference in genome structure between the two *Nasuia* strains. We found complete conservation of synteny in both endosymbiont pairs, reflecting the high level of gene-order stability seen in other ancient endosymbionts (fig. 2) (McCutcheon et al. 2019).

### Higher frequency of SNVs in *Nasuia-MSEV* than in *Sulcia-MSEV*

We then used our new endosymbiont genome assemblies as references for mapping duplex sequence data. We detected a 106-fold higher SNV frequency in *Nasuia-MSEV* (2.18×10^− 5^) than in *Sulcia-MSEV* (2.04×10^−7^) (fig. 3A, supplementary data S1). For both endosymbionts, observed SNV frequencies are above the noise threshold of 2×10^−8^ errors per bp that we have previously established for our duplex protocol (Wu et al. 2020). All 17 of the SNVs identified in *Sulcia-MSEV* were ‘singletons’ (captured by only a single DCS). In contrast, 32 of the 126 positions with SNVs in *Nasuia-MSEV* were detected at ‘high frequency’ (captured by more than one DCS). Accordingly, the 106-fold higher SNV frequency in *Nasuia-MSEV* is partially driven by variants present at high frequency in the experimental populations. We therefore performed a comparison based on independent SNV events (sites in the genome with a SNV) and found a 24-fold higher independent event SNV frequency in *Nasuia-MSEV* than in *Sulcia-MSEV* after normalizing for sequence coverage (fig. 3b, supplementary table 4).

**Fig. 3.**
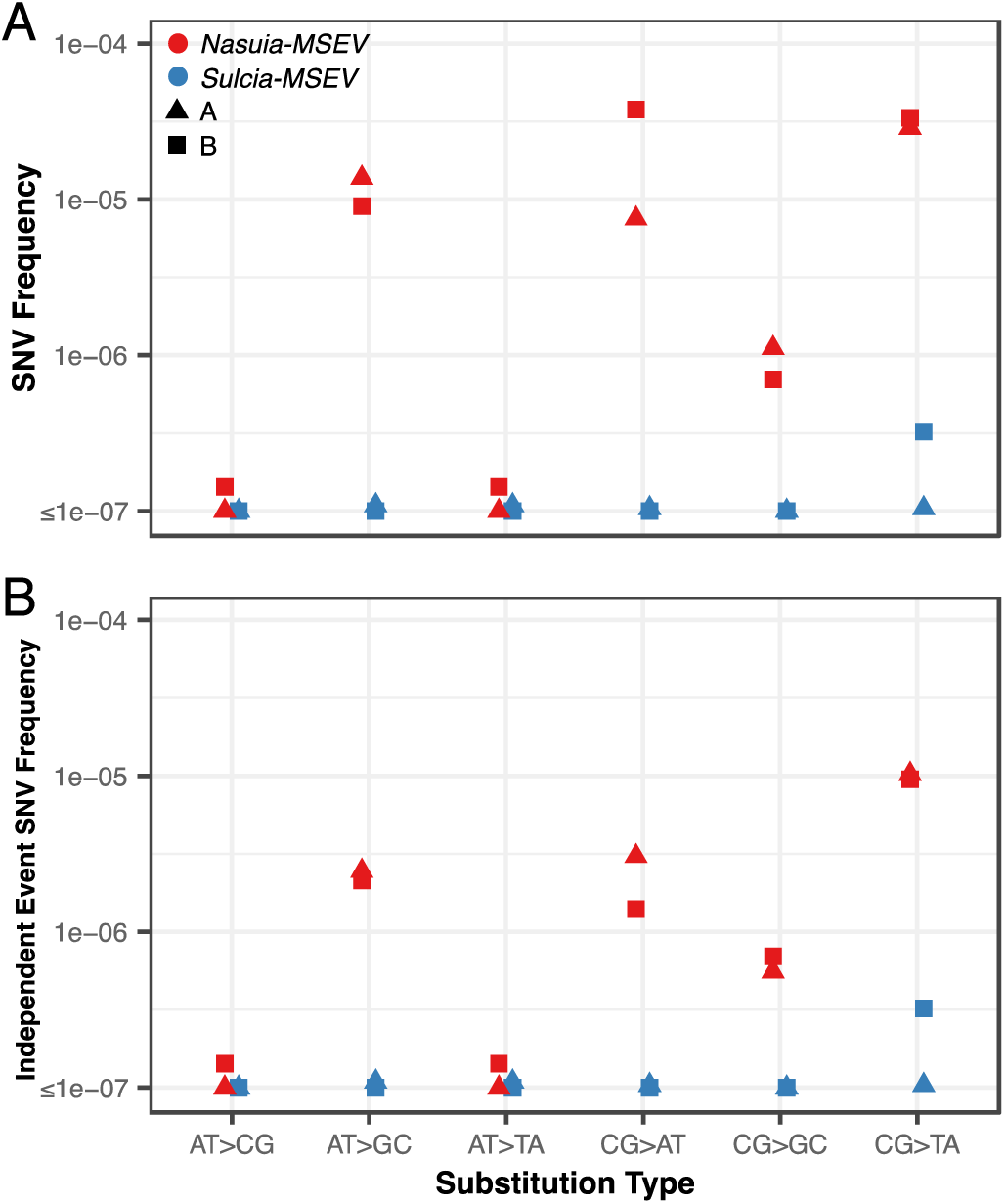
Duplex sequencing variant frequency in *Nasuia-MSEV* and *Sulcia-MSEV*. Variant frequencies are shown **(A)** in terms of total SNV frequency (DCS variants/DCS coverage) and (**B)** in terms of independent event frequency (sites with a variant/DCS coverage). In both, the six mutation types and are normalized by DCS coverage of the corresponding reference bases (for example the AT CG changes are normalized by total AT coverage). *Nasuia-MSEV* is red and *Sulcia-MSEV* is blue. Triangles and squares are experimental populations A and B, respectively. For *Nasuia-MSEV*, the average (of the two experimental populations) independent event frequency (4.97×10^−6^) is reduced relative to the average SNV Frequency (2.18×10^−5^) due to the presence of 32 sites where multiple DCS reads mapped with a variant. There is no difference in *Sulcia-MSEV* variant frequencies (2.04×10^−7^) because all *Sulcia-MSEV* SNVs are singletons (see also supp Fig 1).

For both *Sulcia-MSEV* and *Nasuia-MSEV*, the SNV type with the highest independent event frequency was CG TA transitions, though SNV type comparisons are low powered in *Sulcia-MSEV* due to the small number of variants (supplementary figure S1). In *Nasuia*, AT GC → Transitions and CG AT transversions were also detected at relatively high frequencies. Given the extreme AT bias of the genomes (*Nasuia-MSEV*: 83.1% AT, *Sulcia-MSEV*: 76% AT), we tested if genomic AT content is at equilibrium, in which case the number of AT-increasing mutations would equal the number of AT-decreasing mutations. Of the 126 (independent event) SNVs detected in *Nasuia-MSEV*, significantly less than half (only 43) decreased AT content, while 75 mutations increased AT content (eight changes were AT-neutral) (binomial test, *p* = 0.0004). The five AT-decreasing and six AT-increasing SNVs we detected in *Sulcia-MSEV* provide little power for testing for AT content equilibrium, but do not deviate significantly from equilibrium expectations (binomial test, *p* = 1.0). The proportion of transitions out of all independent SNVs was 0.82 and 0.52 for *Nasuia-MSEV* and *Sulcia-MSEV*, respectively.

Interestingly, nine of the 32 high-frequency SNVs in *Nasuia-MSEV* were detected in both experimental populations A and B (fig. 4). Shared SNVs could have originated in the ancestor of the A and B foundresses and remained polymorphic during the five months that the experimental populations were maintained. Alternatively, the shared SNVs could have arisen independently in the two experimental lines (i.e. homoplasy). There were no SNVs shared across any replicates in *Sulcia-MSEV*.

**Fig. 4.**
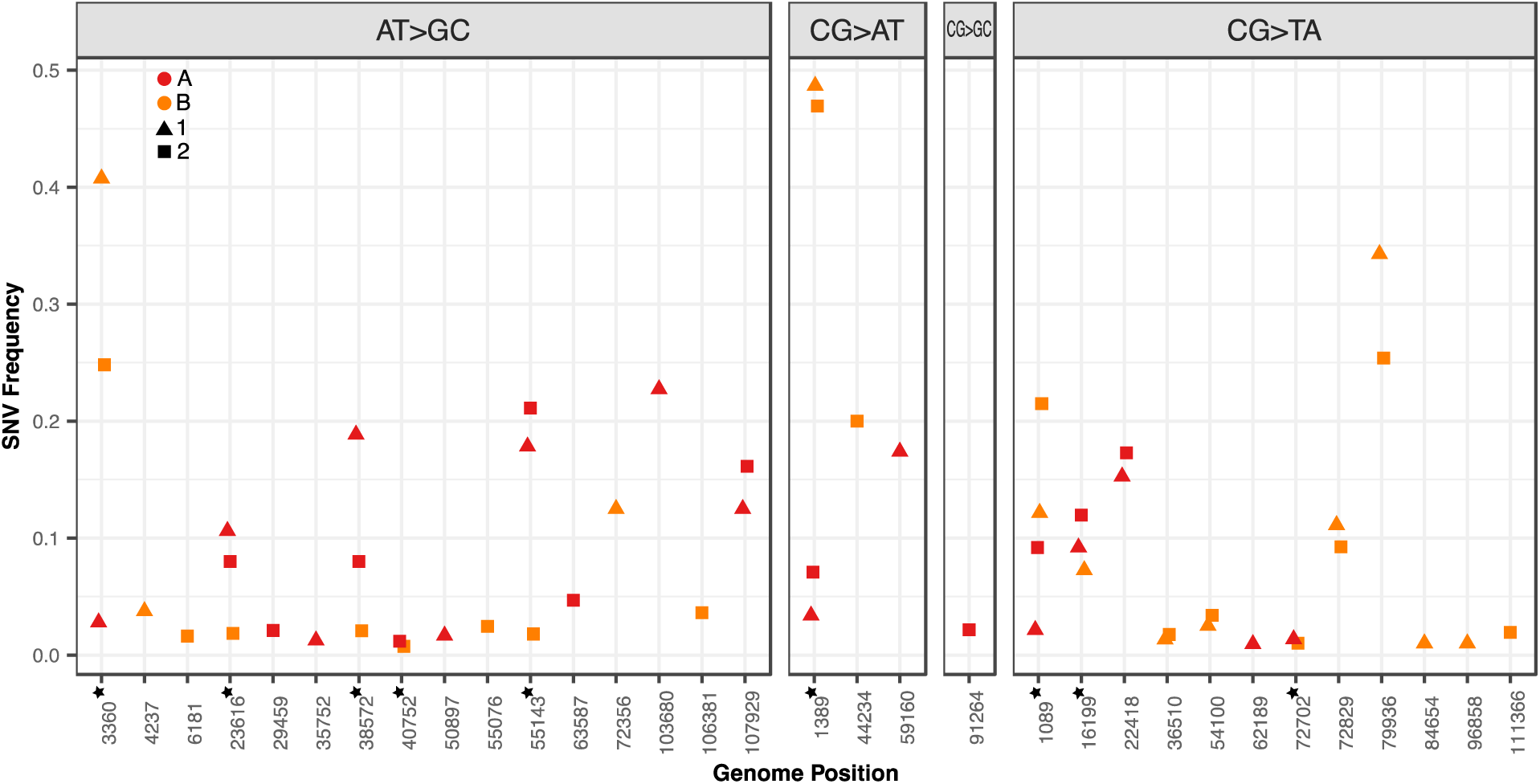
High-frequency variants in *Nasuia-MSEV*. SNV frequencies of the 32 positions in the *Nasuia-MSEV* genome captured by more than one DCS. Frequencies for experimental populations A and B are shown in red and orange, respectively. Experimental populations were subdivided into replicates 1 and 2, which are shown in triangles and squares, respectively. At nine positions in the genome, variants are shared between experimental populations A and B.

In *Nasuia-MSEV* protein-coding regions, we found that the proportion of nonsynonymous changes was significantly reduced in high-frequency SNVs compared to singletons (Fisher’s Exact Test, *p* = 0.0003). Reduction in the proportion of nonsynonymous SNVs in high-frequency SNVs (compared to singletons) occurs in all six substitution types, though only CG → TA transitions are significant on their own (Fisher’s Exact Test, *p* = 0.0179; supplementary figure S2, supplementary table 5). The nucleotide substitution spectra also differ between singletons and high-frequency SNVs. Unlike the bias towards AT-increasing mutations in the overall dataset, the number of high-frequency SNVs that decrease AT content is nearly equivalent to the number that increase AT content (16 and 15, respectively). We could not perform the same analysis for *Sulcia-MSEV* SNVs, as they were all singletons. For both endosymbionts, SNVs are not differentially distributed across intergenic, protein-coding, rRNA, or tRNA regions of the genome (supplementary figure S3, supplementary table 6).

### No strand-specific mutation asymmetry detected in *Nasuia-MSEV* protein-coding genes

Coding strands in CDS regions of the *Nasuia-MSEV* genome are enriched for G compared to C (positive GC skew of 0.16 or 1.4 Gs for every C). In endosymbiotic genomes, positive GC skew on the leading strand of DNA replication is thought to result from asymmetrical C → T changes on the leading strand, which is more susceptible to cytosine deamination due to its prolonged existence in a single-stranded state (Klasson and Andersson 2006). During transcription, the increased single-stranded exposure of the coding DNA strand has also been proposed to lead to asymmetrical C → T changes in bacterial genomes (Francino and Ochman 1997).

We tested if mutation drives GC skew on coding stands of *Nasuia-MSEV* by comparing reciprocal mutations (C → T vs. G → A) to expected counts given C and G coverage. We did not find evidence for asymmetry in reciprocal mutations in CG → TA transitions, or for any of other five substitution types in *Nasuia-MSEV* coding strands in CDS regions. (binomial test, all p-values ≥ 0.16) (supplementary table 7). We did not attempt to perform the reciprocal strand analysis for *Sulcia-MSEV* given the small amount of SNVs.

### Comparison of indels in *Nasuia-MSEV* and *Sulcia-MSEV*

Overall, average indel frequencies for *Nasuia-MSEV* and *Sulcia-MSEV* are similar (4.92×10^−6^ and 4.51×10^−6^ respectively). *Nasuia-MSEV* has a higher frequency of deletions than insertions while *Sulcia-MSEV* has a higher frequency of insertions than deletion s (fig. 5a, supplementary data S2). However, it is important to recognize that during genome assembly, determination of the dominant repeat number in regions with repeat length heterogeneity can influence whether an indel is considered a deletion or an insertion, so comparisons of the frequencies of deletions vs. insertions should be cautiously interpreted. Indeed, when the three hypervariable regions that were identified during genome assembly (see Materials and Methods) are excluded from *Sulcia* indel analysis, insertions become less frequent than deletions (supplementary table 8). For all downstream analysis, no regions were excluded (i.e. the full data set was considered).

**Fig. 5.**
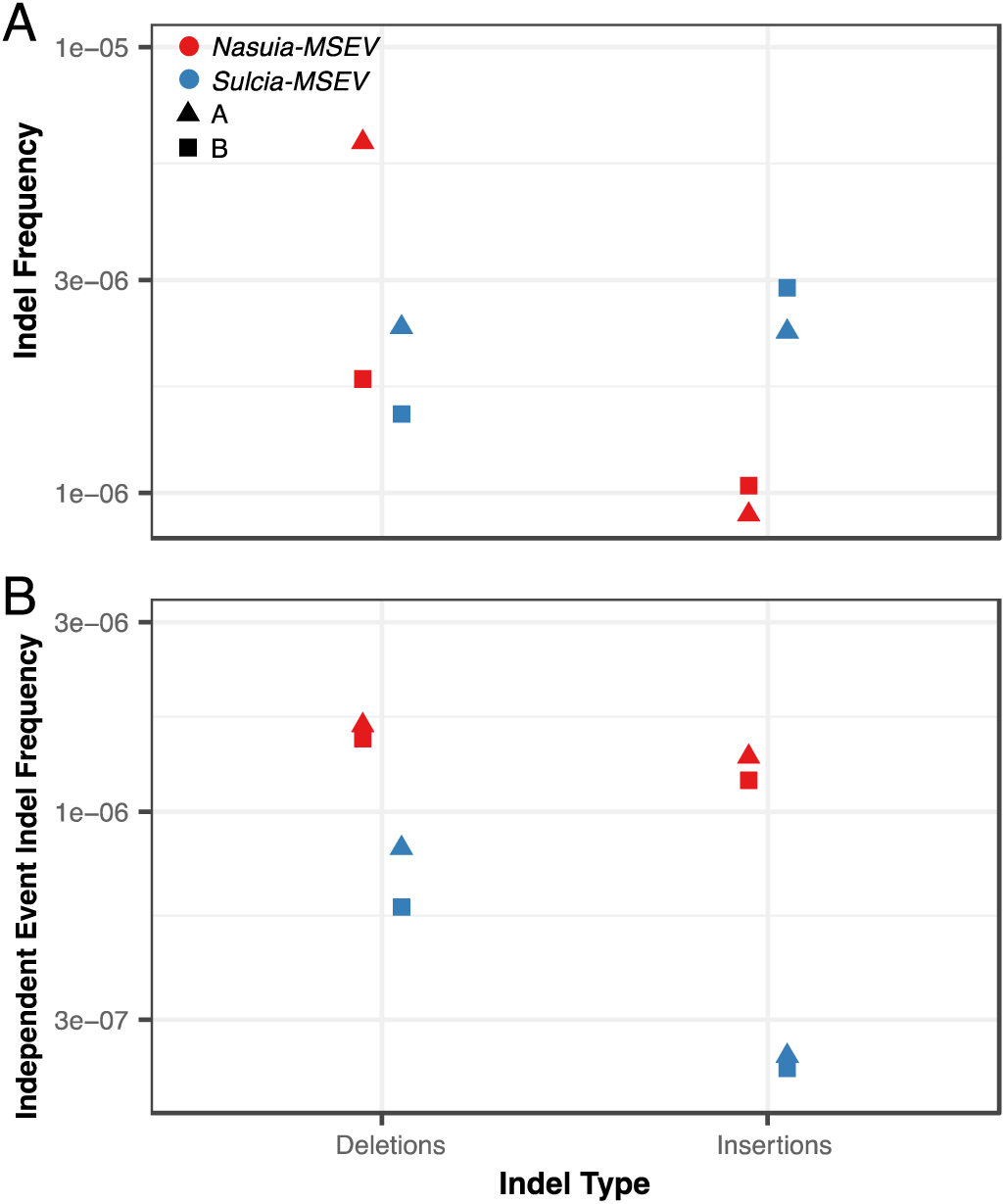
Indels frequencies in *Nasuia-MSEV* and *Sulcia-MSEV*. Indel frequencies shown in terms of **(A)** total frequency (DCS indels/DCS coverage) and **(B)** Independent event frequency (sites with an indel/DCS coverage). Overall average indel frequencies for *Nasuia-MSEV* and *Sulcia-MSEV* are similar – but deletions are more common in *Nasuia-MSEV* and insertions are more common in *Sulcia-MSEV*. Independent event frequencies are higher for *Nasuia-MSEV* than for *Sulcia-MSEV. Nasuia-MSEV* is red and *Sulcia-MSEV* is blue. Triangles and squares are experimental populations A and B, respectively.

Both endosymbionts have indels that are present at high frequencies, which are sometimes shared across experimental populations A and B. Like with high-frequency SNVs in *Nasuia-MSEV*, these indels could arise from homoplasy or from shared ancestry and persistence in a polymorphic state during the growth of experimental cultures. Local sequence context plays a large role in indel distribution across both genomes as 94.1% and 98.4% of indels overlap with repetitive genomic sequence (microsatellites and homopolymers) for *Nasuia-MSEV* and *Sulcia-MSEV*, respectively. Given the apparent influence of local sequence context in indel occurrence, and the lack of local sequence effect on SNV occurrence, we suspect that high frequency indels often reflect multiple homoplasious events (e.g., long homopolymers can be subject to “multiple hits”). Our indel analysis therefor emphasizes total indel frequency rather than independent event indel frequency (fig. 5b).

We found that *Nasuia-MSEV* has a higher frequency of homopolymer-associated indels, while *Sulcia-MSEV* has a higher frequency of microsatellite-associated indels (fig. 6a). These differences do not merely reflect a difference in the repetitive landscape of the symbiont genomes, as the frequencies are normalized by the specific coverage of a given repeat type (microsatellite or homopolymer), and *Nasuia-MSEV* contains more of both repeat type per kb than *Sulcia-MSEV* (supplementary table 1). For both endosymbionts, we find that as homopolymers increase in length, indel frequency also increases (fig. 6b).

**Fig. 6.**
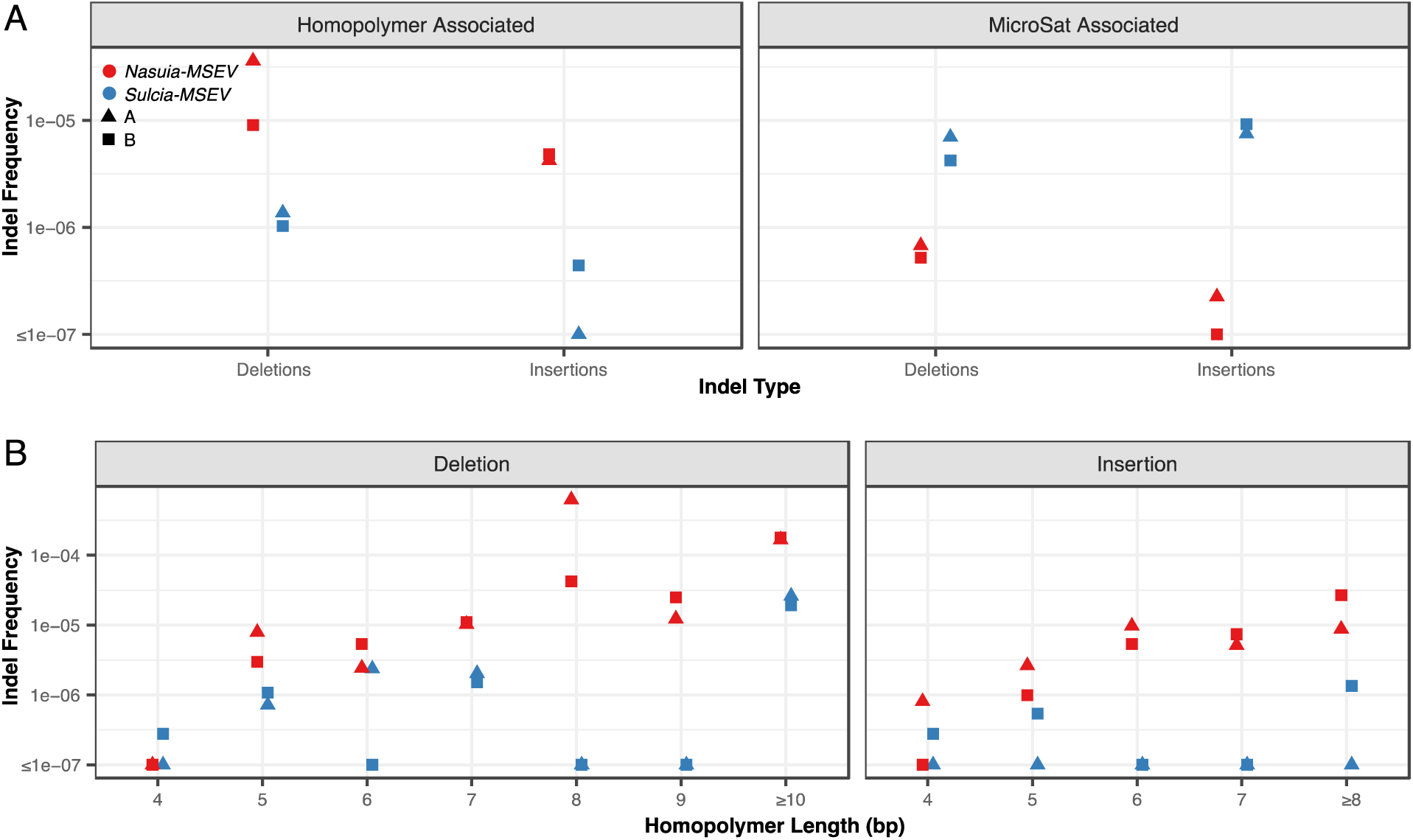
Frequency of repeat-associated indels in *Nasuia-MSEV* and *Sulcia-MSEV*. **(A)** Frequencies of indels associated with homopolymers and microsatellites and **(B)** frequencies of homopolymer-associated indels as function of homopolymer length. Frequencies are calculated as the total number of DCS indels in a repeat category divided by DCS coverage of that repeat category. 94.1% of all *Nasuia-MSEV* indels and 98.4% of all *Sulcia-MSEV* indels overlap with homopolymers or microsatellites. 90.9 % and 96.1 % of all indels are expansions or contractions of existing repeats for *Nasuia* and *Sulcia*, respectively. For *Nasuia-MSEV*, the majority of repeat-associated indels are located in homopolymers, and deletions are more common than insertions. For *Sulcia-MSEV*, the majority of indels are located in microsatellites. For both endosymbionts, the frequency of insertions and deletions increase as homopolymer length increases. *Nasuia-MSEV* is red, and *Sulcia-MSEV* is blue. Triangles and squares are experimental populations A and B, respectively.

## DISCUSSION

### Rapid evolution of *BetaSymb* likely driven by higher mutation rate

*BetaSymb* (represented by *Nasuia* in leafhoppers and other names depending on the insect host; *Vidani* in planthoppers and *Zinderia* in spittlebugs) evolves much more rapidly that *Sulcia*, despite the partner endosymbionts having occupied a nearly identical ecological niche (bacteriocytes in a shared host) for ∼300 million years (Bennett, Abbà, et al. 2016; Bennett and Mao 2018). In addition, *BetaSymb* experiences increased evolutionary turnover and has been replaced in at least six auchenorrhynchan lineages (Bennett and Moran 2015). This accelerated evolution with respect to *Sulcia* does not appear to be limited to *BetaSymb*, as some of its lineage-specific replacements such as *Hodgkinia* (Alphaproteobacteria) and *Baumannia* (Gammaproteobacteria) show similar asymmetries (Bennett, McCutcheon, et al. 2016; Campbell et al. 2017). Our finding of a higher frequency of *de novo* SNVs in *Nasuia-MSEV* (fig. 3) suggests the increased evolutionary rate in *BetaSymb* is likely driven by mutation.

Because variant frequencies are the product of both the mutation rate (μ) and the effective population size (N_e_), it is important to recognize that a higher N_e_ in *Nasuia-MSEV* could also lead to the 24 to 106-fold higher SNV frequency we observed in *Nasuia-MSEV*. However, current evidence does not support a larger N_e_ for *Nasuia*. Sequence coverage comparisons from our bacteriome shotgun library reveal a 1.5-fold higher per bp depth of coverage in *Sulcia-MSEV* than in *Nasuia-MSEV* (Pedersen and Quinlan 2018). This suggests the standing population size may actually be smaller in *Nasuia-MSEV* than in *Sulcia-MSEV*, though bias against AT-rich sequences during library amplification could also drive lower coverage of the *Nasuia-MSEV* genome. The standing population size does not account for the possibility that there may be differences in the intensity of the transmission bottlenecks for the two endosymbionts. Microscopy has revealed that, while *Sulcia* and *Nasuia* are housed in distinct bacteriocyte types, they both successfully migrate to and infect terminal oocytes (Szklarzewicz et al. 2016). It is not known, however, if the number of transmitted bacteria differs for *Nasuia* vs. *Sulcia*. Finally, a larger N_e_ in *Nasuia-MSEV* should lead to more efficient purifying selection, which would be inconsistent with the fact that *Nasuia* exhibits more extensive genome decay (high AT content, reduced size, etc.) over phylogenetic scales.

Interestingly, 32 variants in *Nasuia-MSEV* were present in ‘high frequency’ (detected in more than one DCS). Compared to singletons, high-frequency SNVs in *Nasuia-MSEV* are less likely to result in nonsynonymous changes (supplementary figure 2), suggesting they have been filtered by selection. This idea is further supported by the observation that high frequency SNVs are in equilibrium with genomic AT levels, while singletons display an AT mutation bias. Singletons in our data thus most accurately represent the true mutation spectra, before it can be shaped by selection. As such, despite the already extreme AT-richness of the *Nasuia* genome, it appears that mutation bias would further erode GC content in the absence of selection (Hershberg and Petrov 2010; Hildebrand et al. 2010; Long, Sung, et al. 2018). Our observation of rapid filtering of high frequency events in *Nasuia-MSEV* raises the possibility that a previous study finding no AT mutation bias in *Buchnera aphidicola* (an ancient endosymbiont of aphids) (Moran et al. 2009) likely did so based on a set of mutations that had been subject to selection. While our data demonstrate that selection is able to maintain some level of *Nasuia* sequence conservation, it remains possible that *Nasuia* and *Sulcia* are differentially constrained.

Nine of the 32 *Nasuia-MSEV* high frequency SNVs were shared between experimental populations A and B, which were reproductively isolated for six months (approximately six host generations; Capinera 2008). While it is possible that shared SNVs resulted from homoplasy, some shared SNVs have risen to very high frequency (greater than 45% in one case), which demonstrates they are at least old enough to have spread to a large portion of the experimental population. We find it most likely that shared SNVs were ancestral to endosymbiont populations present in the initial insect foundresses and persisted as polymorphisms through the duration of insect growth.

### Can differences in DNA replication and repair machinery explain greater genome conservation in *Sulcia*?

While *Nasuia-MSEV* and *Sulcia-MSEV* lack most subunits of the primary bacterial polymerase (DNA Pol III), they both retain α and ε subunits (*dnaE* and *dnaQ*), which in *Escherichia coli* perform polymerization and 3′ → 5′ exonuclease activity, respectively (Fijalkowska et al. 2012). *Nasuia-MSEV* also uniquely retains the β subunit (*dnaN*), which facilitates binding between subunit α and the template DNA strand (Gui et al. 2011). *Nasuia-MSEV* retains two additional DNA replication and repair (RR) genes lost in *Sulcia-MSEV*: *dnaB* (helicase) and *ssb* (single stranded binding protein) (Bennett and Moran 2013).

Only one RR gene, *mutS*, is absent in *Nasuia-MSEV*, but retained in *Sulcia-MSEV*. In other bacteria, the MutS protein recognizes and initiates repair of mismatched bases and small indels (Dettman et al. 2016; Long, Miller, et al. 2018). While the role of *mutS* in *Sulcia* has never been experimentally validated, it contains all five of the domains present in the *E. coli mutS* (Ogata et al. 2011; Groothuizen and Sixma 2016). The extremely low frequency of SNVs we observed in *Sulcia-MSEV* (fig 1, Supp Table 4) could result from *mutS*-directed mismatch repair (MMR).

Mutation accumulation experiments with *Pseudomonas aeruginosa* reveal *mutS* mutants experience a 230-fold increase in indels (average length of 1.1 bp) compared to WT lines, demonstrating the importance of *mutS* in short indel surveillance (Dettman et al. 2016). While we observed a high frequency of indels in *Sulcia-MSEV*, most were expansions or contractions of repeats at existing microsatellites. In fact, only 19 of the 385 indel containing DCS reads in *Sulcia-MSEV* were of 1 bp in length (compared to 145 of 154 in *Nasuia-MSEV*). Even more striking is the complete lack of 1-bp indels in the alignment of *Sulcia-MSEV* with *Sulcia* from *M. quadrilineatus* (compared to 48 1-bp indels in the equivalent *Nasuia* alignment) (supplementary table 1). The rarity of 1-bp indels in *Sulcia-MSEV* thus further supports the idea that *mutS* is critical for maintaining *Sulcia* genome conservation through surveillance of homopolymer expansions/contractions.

Interestingly, the genome of *Sulcia* in leafhoppers actually has a reduced set of RR genes compared to those found in *Sulcia* genomes in other auchenorrhynchan lineages. *Sulcia-CARI*, an endosymbiont of spittlebugs, possesses the two DNA Pol III subunits found in *Sulcia-MSEV*, but has also retained in subunits *dnaB, dnaN, dnaX*, and polymerase accessory subunits *holA* and *holB*. Additional RR genes retained by *Sulcia-CARI*, but absent in leafhopper *Sulcia* genomes, include gyrase genes *gryA, gryB* and MMR genes *mutL*, and *mutH* (McCutcheon and Moran 2010; Bennett and Moran 2013). Despite lacking the above listed RR genes, leafhopper *Sulcia* genomes remain remarkably stable, as demonstrated by phylogenetic comparisons at multiple levels of leafhopper divergence in this and previous studies (Bennett, Abbà, et al. 2016; Mao et al. 2017).

Given that *mutS* works in concert with *mutL* and *mutH* in typical bacterial MMR, it is curious that *mutH* has been completely lost and *mutL* has been pseudogenized in leafhopper *Sulcia* genomes (Bennett and Moran 2013; Bennett, Abbà, et al. 2016). A recent assay of *M. quadrilineatus* gene expression revealed that in the bacteriome, leafhopper hosts often overexpresses homologs of RR genes that are absent from endosymbiont genomes, raising the possibility that leafhopper hosts may support endosymbiont genome replication and repair (Mao et al. 2018). Host support has been hypothesized to enable endosymbiont gene loss (Hansen and Moran 2011; Husnik et al. 2013; Russell et al. 2013; Sloan et al. 2014), and a recent study has demonstrated that the partner endosymbionts of cicadas likely rely of host encoded machinery to acquire correctly processed tRNAs (Van Leuven et al. 2019). However, none of the four putative *mutL/mutH* homologs in the *M. quadrilineatus* transcriptome showed substantial overexpression in the *Sulcia* specific bacteriocyte (compared to expression in the insect body) (supplementary table 9). Thus, it remains unclear if *Sulcia mutS* is participating in conventional MMR or if it has acquired novel functions in support of *Sulcia* genome conservation. Functional characterization of MMR in leafhopper *Sulcia* would provide valuable insight into what appears to be an atypical role for an ancient and highly conserved mismatch recognition protein (Ogata et al. 2011; Jiricny 2013).

The missing *mutS* in *Nasuia* provides an additional opportunity for host support. Mao et. al. (2018) reported strong overexpression of one host *mutS* homolog (referred to as TRINITY_DN66078_c1_g1 in that study; see supplementary table 9) in the *Nasuia* bacteriome, but it is highly unlikely that this is involved in MMR for two reasons. First, the identified homolog is related to eukaryotic *MSH4*, which is a gene family that lacks a mismatch recognition domain and is involved in recombination rather than MMR (Ogata et al. 2011). Furthermore, it appears to encode only a partial MSH4 protein fused with a C-terminal sequence of unidentified origin. Therefore, it is likely that *Nasuia* lacks conventional MMR function.

While *mutS* may play a role in *Sulcia* genome conservation, retention of *mutS* is not a cure-all. For example, *BetaSymb* in spittlebugs (named *Candidatus* Zinderia insecticola) retains the entire mismatch repair gene set (*mutS, mutL*, and *mutH*) but displays a rate of evolution typical of *Nasuia* and other *BetaSymb* lineages (Bennett and Moran 2013; Koga et al. 2013). We posit that the *mutS* in *Sulcia* may be particularly efficient at recognizing and removing mismatched bases and short indels. All *Sulcia* genomes analyzed to date retain *mutS*, but identification of a *Sulcia* lineage that has lost *mutS* would facilitate a test of the hypothesis that *mutS* may be the key to *Sulcia* genome conservation.

## CONCLUSION

The two ancestral endosymbionts of auchenorrhynchan insects vary dramatically in their propensity for extinction vs. retention, thus providing a natural comparative framework for investigating what facilitates long-term, stable endosymbiotic relationships. Compared to *Sulcia*, which exhibits remarkable genome conservation for an ancient endosymbiont and has been nearly universally retained, *BetaSymb* displays substantially elevated rates of molecular evolution. We have shown that *Nasuia-MSEV*, a representative of *BetaSymb* from leafhoppers, has a substantially higher frequency of *de novo* mutations than *Sulcia-MSEV*. Our analysis supports the hypothesis that the evolutionary rate variation in *Sulcia* vs. *BetaSymb* is driven by differences in mutational input. We posit that the low mutation rate in *Sulcia* may also explain why *Sulcia* is so consistently retained in diverse auchenorrhynchan lineages, while partner endosymbionts are apparently much more transient.

## DATA AVAILABILITY

The raw sequencing reads are available via the NCBI Sequence Read Archive under accessions SRR12112868, SRR12112867, SRR12112866, SRR12112865, SRX8635374 and SRR12112864. Genome annotations for *Nasuia-MSEV* and *Sulcia-MSEV* are available via GenBank under accessions SAMN15504451 and SAMN15504450, respectively.

## ACKNOWEDGEMENTS

We thank Meng Mao for providing Trinity assemblies and differential expression data. We also thank Kristen Poff at UH Manoa for early help with insect rearing. This work was supported by the National Institutes of Health (R01 GM118046) and a National Science Foundation graduate fellowship (DGE 1450032).

## SUPPLEMENTAL FIGURES

**Supplemental Fig. 1.**
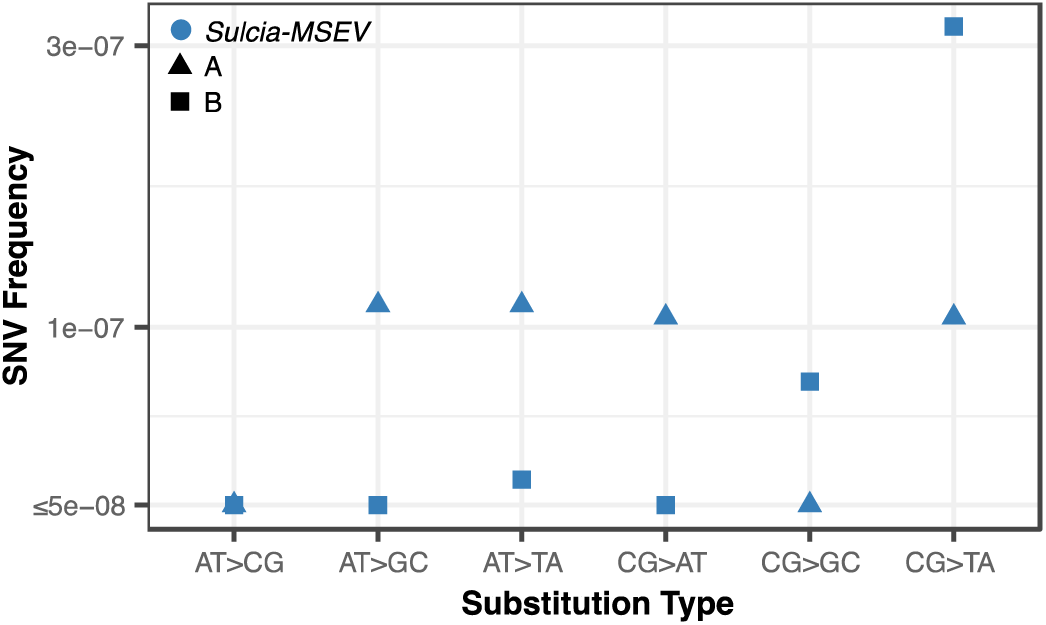
SNVs frequency in *Sulcia-MSEV*. CG → TA transitions are the most common type in population B. In population A there is little variation in frequency between AT → GC, CG → TA, AT → TA, and CG → AT substitutions.

**Supplemental Fig. 2.**
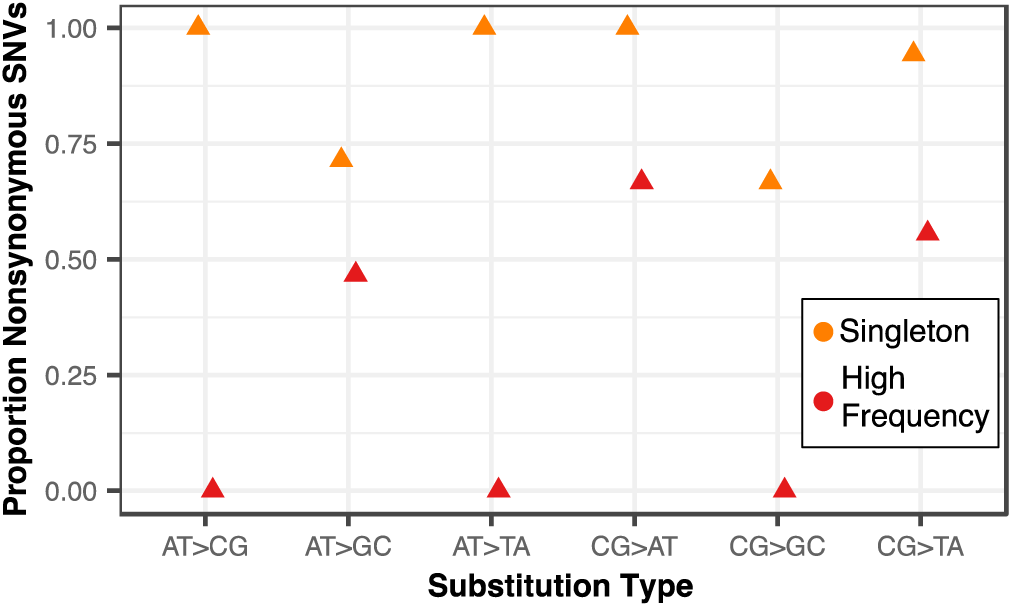
Proportion nonsynonymous SNVs in *Nasuia-MSEV* CDS regions. The proportion of nonsynonymous SNVs is calculated as number of SNVs/total SNVs. In all substitution types, the proportion of nonsynonymous SNPs is reduced in the high-frequency class compared to the singleton.

**Supplemental Fig. 3.**
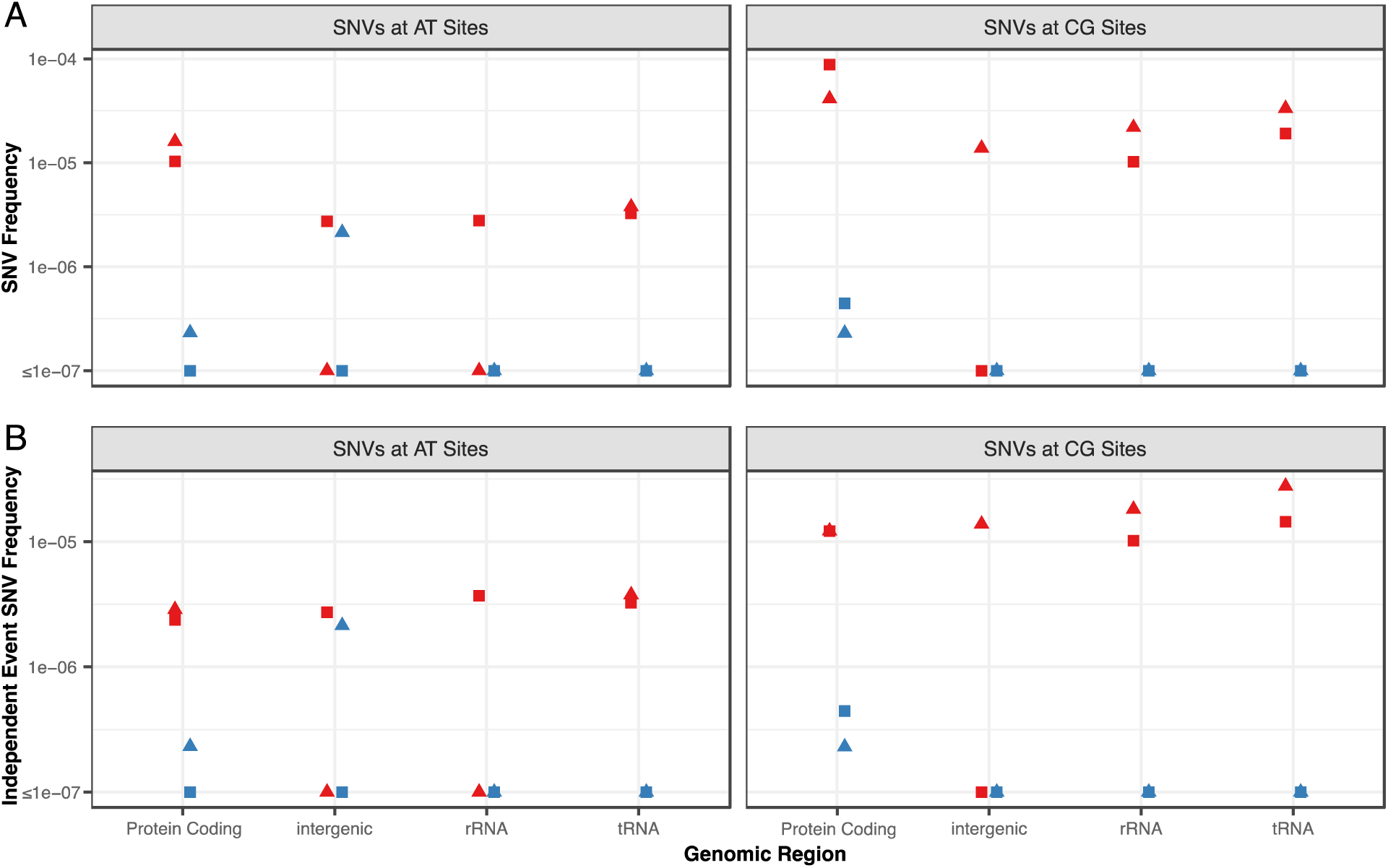
SNV frequencies by region for AT sites and CG sites in *Nasuia* and *Sulcia*. For regions of the genome with different functional annotations, variant frequencies are shown **(A)** in terms of total SNV frequency (DCS variants per region/DCS coverage region) and (**B)** in terms of independent event frequency (sites with a variant per region/DCS coverage per region). For both endosymbionts, independent event frequencies are distributed evenly across regions of the genome (binomial test, all p-values ≥ 0.076. supplementary table 6).

## SUPPLEMENTAL TABLES

**Supplementary Table 1.** Repetitive sequence content of the Nasuia and Sulcia genomes.

|  | Homopolymer Counts |  | Microsatellite Counts |  |
| --- | --- | --- | --- | --- |
|  | 4 + bp in length | 6 + bp in length | 4 + bp in length | 6 + bp in length |
| *Nasuia* | 6283 | 1817 | 8452 | 4534 |
| *Sulcia* | 7092 | 1791 | 12196 | 6043 |

**Supplementary Table 2.** DCS coverage and percent mapping for 4 reps. For the four duplex libraries (A1, A2, B1, B2), an average of 18.45% of DCS reads mapped to one of the endosymbiont genomes, resulting in an average per bp coverage of 71.2 and 112.1 for *Nasuia* and *Sulcia*, respectively. The majority of non-mapping reads are presumably derived from host (*M. sp. nr. severini*) DNA, which is expected to be common as the duplex libraries were created from DNA isolated from *M. sp. nr. severini* bacteriome tissue.

| Replicate | Total DCS bp | Percent Mapping | Total *Nasuia* Coverage | *Nasuia* per bp coverage | Total *Sulcia* Coverage | *Sulcia* per bp coverage |
| --- | --- | --- | --- | --- | --- | --- |
| A1 | 173374467 | 18.38 | 9498361 | 83.53 | 22365568 | 117.20 |
| A2 | 149941788 | 13.13 | 5051784 | 44.42 | 14642073 | 76.73 |
| B1 | 154509864 | 19.67 | 7910980 | 69.57 | 22480070 | 117.80 |
| B2 | 161221215 | 22.65 | 10411542 | 91.56 | 26101409 | 136.78 |

**Supplementary Table 3.** Indel lengths and counts from alignments of endosymbionts from. *M. quadrilineatus* and *M sp. nr. severini*. Indel counts were tabulated with a custom script.

| Sulcia Pairs |  | Nasuia Pairs |  |
| --- | --- | --- | --- |
| Indel Length (bp) | Count | Indel Length (bp) | Count |
| 3 | 1 | 1 | 48 |
| 5 | 4 | 2 | 12 |
| 6 | 26 | 3 | 35 |
| 7 | 1 | 4 | 1 |
| 9 | 3 | 5 | 6 |
| 10 | 1 | 6 | 4 |
| 12 | 6 | 7 | 1 |
| 18 | 3 | 9 | 1 |
| 21 | 1 | 11 | 1 |
| 24 | 2 | 50 | 1 |
| 28 | 1 | 60 | 1 |
| 42 | 1 | 65 | 1 |
| 48 | 1 | 71 | 1 |
| 50 | 1 | 1352 | 1 |
| 54 | 1 |  |  |
| 60 | 1 |  |  |

**Supplementary Table 4.** SNV frequency and independent event frequency for the four replicates. Substitution types for both SNV frequency and independent event SNV frequency are normalized based on genome specific reference base coverage (i.e. AT → CG, AT → GC and AT → TA types are all normalized by genome specific duplex AT coverage).

| Replicate | Substitution Type | *Nasuia-MSEV* |  | *Sulcia-MSEV* |  |
| --- | --- | --- | --- | --- | --- |
|  |  | SNV frequency | independent event SNV frequency | SNV frequency | independent event SNV frequency |
| A1 | AT>CG | 0 | 0 | 6.02E-08 | 6.02E-08 |
| A1 | AT>GC | 1.81E-06 | 1.37E-05 | 1.20E-07 | 1.20E-07 |
| A1 | AT>TA | 1.39E-07 | 1.39E-07 | 0 | 0 |
| A1 | CG>AT | 2.18E-06 | 5.66E-06 | 1.74E-07 | 1.74E-07 |
| A1 | CG>GC | 4.35E-07 | 4.35E-07 | 0 | 0 |
| A1 | CG>TA | 1.26E-05 | 2.83E-05 | 1.74E-07 | 1.74E-07 |
| A1 | Total | 5.16E-06 | 1.88E-05 | 2.24E-07 | 2.24E-07 |
| A2 | AT>CG | 0 | 0 | 0 | 0 |
| A2 | AT>GC | 3.73E-06 | 1.36E-05 | 9.26E-08 | 9.26E-08 |
| A2 | AT>TA | 0 | 0 | 2.78E-07 | 2.78E-07 |
| A2 | CG>AT | 4.63E-06 | 1.08E-05 | 0 | 0 |
| A2 | CG>GC | 7.71E-07 | 2.31E-06 | 0 | 0 |
| A2 | CG>TA | 6.17E-06 | 2.93E-05 | 0 | 0 |
| A2 | Total | 5.74E-06 | 2.10E-05 | 2.73E-07 | 2.73E-07 |
| B1 | AT>CG | 1.67E-07 | 1.67E-07 | 0 | 0 |
| B1 | AT>GC | 1.84E-06 | 1.00E-05 | 0 | 0 |
| B1 | AT>TA | 0 | 0 | 1.20E-07 | 1.20E-07 |
| B1 | CG>AT | 1.55E-06 | 3.88E-05 | 0 | 0 |
| B1 | CG>GC | 1.04E-06 | 1.04E-06 | 0 | 0 |
| B1 | CG>TA | 1.09E-05 | 3.68E-05 | 5.17E-07 | 5.17E-07 |
| B1 | Total | 4.80E-06 | 2.64E-05 | 2.22E-07 | 2.22E-07 |
| B2 | AT>CG | 1.24E-07 | 1.24E-07 | 0 | 0 |
| B2 | AT>GC | 2.36E-06 | 8.33E-06 | 5.12E-08 | 5.12E-08 |
| B2 | AT>TA | 2.49E-07 | 2.49E-07 | 0 | 0 |
| B2 | CG>AT | 1.27E-06 | 3.68E-05 | 0 | 0 |
| B2 | CG>GC | 4.23E-07 | 4.23E-07 | 1.52E-07 | 1.52E-07 |
| B2 | CG>TA | 8.46E-06 | 3.05E-05 | 1.52E-07 | 1.52E-07 |
| B2 | Total | 4.42E-06 | 2.21E-05 | 1.15E-07 | 1.15E-07 |

**Supplementary Table 5.**
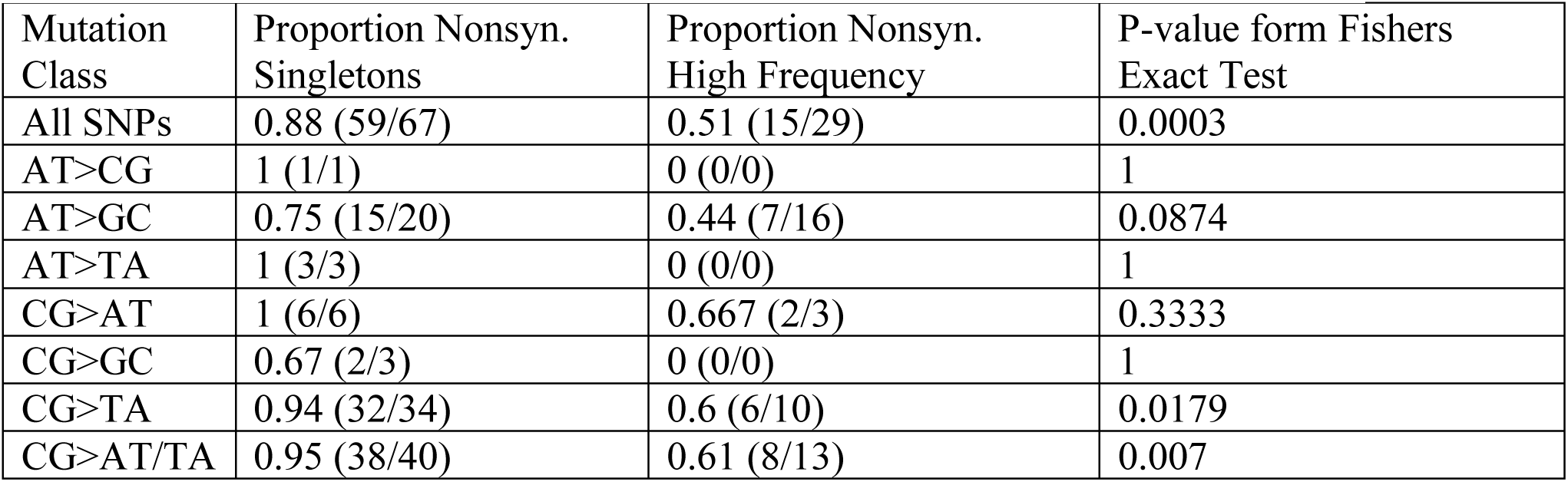
Comparison of the proportion of nonsynonymous SNVs in *Nasuia-MSEV* CDS regions between high-frequency and singleton SNVs.

| Mutation Class | Proportion Nonsyn. Singletons | Proportion Nonsyn. High Frequency | P-value form Fishers Exact Test |
| --- | --- | --- | --- |
| All SNPs | 0.88 (59/67) | 0.51 (15/29) | 0.0003 |
| AT>CG | 1 (1/1) | 0 (0/0) | 1 |
| AT>GC | 0.75 (15/20) | 0.44 (7/16) | 0.0874 |
| AT>TA | 1 (3/3) | 0 (0/0) | 1 |
| CG>AT | 1 (6/6) | 0.667 (2/3) | 0.3333 |
| CG>GC | 0.67 (2/3) | 0 (0/0) | 1 |
| CG>TA | 0.94 (32/34) | 0.6 (6/10) | 0.0179 |
| CG>AT/TA | 0.95 (38/40) | 0.61 (8/13) | 0.007 |

**Supplementary Table 6.** Proportion of SNVs compared to proportion of coverage in genomic regions with variants.

| Reference | SNV class | Region | Proportion of coverage | Proportion of SNVs | P-value from the binomial test |
| --- | --- | --- | --- | --- | --- |
| Nasuia | SNVs at AT Sites | CDS | 0.868 | 0.870 | 1.000 |
| Nasuia | SNVs at AT Sites | intergenic | 0.026 | 0.022 | 1.000 |
| Nasuia | SNVs at AT Sites | rRNA | 0.083 | 0.087 | 0.790 |
| Nasuia | SNVs at AT Sites | tRNA | 0.023 | 0.022 | 1.000 |
| Nasuia | SNVs at CG Sites | CDS | 0.785 | 0.700 | 0.076 |
| Nasuia | SNVs at CG Sites | intergenic | 0.021 | 0.013 | 1.000 |
| Nasuia | SNVs at CG Sites | rRNA | 0.145 | 0.200 | 0.153 |
| Nasuia | SNVs at CG Sites | tRNA | 0.049 | 0.088 | 0.115 |
| Sulcia | SNVs at AT Sites | CDS | 0.945 | 0.941 | 1.000 |
| Sulcia | SNVs at AT Sites | intergenic | 0.018 | 0.059 | 1.000 |
| Sulcia | SNVs at CG Sites | CDS | 0.907 | 0.900 | 1.000 |
| Sulcia | SNVs at CG Sites | intergenic | 0.012 | 0.100 | 0.110 |

**Supplementary Table 7.** Analysis of asymmetry in reciprocal mutations for the six substitution types in *Nasuia-MSEV* coding strands (protein-coding regions).

| Substitution Type | Coding Strand Bass | Coverage Proportion | SNV Proportion | P-value from the binomial test |
| --- | --- | --- | --- | --- |
| AT→CG | A | 0.5441 | 1 | 1 |
| AT→GC | A | 0.5441 | 0.583 | 0.7387 |
| AT→TA | A | 0.5441 | 0.66 | 1 |
| CG→AT | C | 0.4043 | 0.22 | 0.328 |
| CG→GC | C | 0.4043 | 0.66 | 0.5696 |
| CG→TA | C | 0.4043 | 0.295 | 0.1672 |

**Supplementary Table 8.** Indels in *Sulcia* with three hypervariable regions excluded.

|  | All sites included |  | 2 hypervariable regions excluded |  |
| --- | --- | --- | --- | --- |
|  | Insertion Frequency | Deletion Frequency | Insertion Frequency | Deletion Frequency |
| A | 2.29E-06 | 2.35E-06 | 1.02E-06 | 1.54E-06 |
| B | 2.88E-06 | 1.50E-06 | 8.23E-07 | 9.05E-07 |
| A and B Average | 2.58E-06 | 1.92E-06 | 9.25E-07 | 1.22E-06 |

**Supplementary Table 9.** Differential expression of genes in the MMR pathway in *Nasuia* and *Sulcia* specific bacteriocytes from Mao et al. 2018. Trinity assemblies were identified through reciprocal blastp against known MMR orthologs in pea aphids (*Acyrthosiphon pisum*) and fruit flies (*Drosophila melanogaster*). See reviews for eukaryotic orthologs of mutS (Ogata et al. 2011), and all MMR proteins (Jiricny 2013).

| Trinity Assembly ID | annotation (reciprocal blastp) | putative bacterial ortholog | Body (FPKM) | *Nasuia* Bacteriocyte (FPKM (FC)) | *Sulcia* Bacteriocyte (FPKM (FC)) |
| --- | --- | --- | --- | --- | --- |
| TRINITY_DN62497_c0_g1 | Msh2 | MutS | 2.00 | 9.9 (4.95) | 6.7 (3.35) |
| TRINITY_DN67078_c0_g1 | Msh6 full | MutS | 9.00 | 1.28 (2.57) | 0.93 (1.88) |
| TRINITY_DN64509_c0_g3 | Msh6 partial | MutS | 1.50 | 6.8 (4.53) | 3.2 (2.13) |
| TRINITY_DN66078_c1_g1 | Msh4-Like | MutS | 0.50 | 27.5 (55) | 5.7 (11.4) |
| TRINITY_DN65597_c0_g2 | Mlh1 | MutL | 1.79 | 2.02 (1.13) | 2.11 (1.18) |
| TRINITY_DN55227_c0_g2 | PMS1 | MutL | 5.08 | 19.11 (3.76) | 10.77 (2.12) |
| TRINITY_DN65716_c0_g1 | PMS2 | MutL/MutH | 1.69 | 2.35 (1.39) | 1.74 (1.03) |
| TRINITY_DN63885_c0_g1 | EXO1 | MutH | 1.53 | 8.06 (5.26) | 5.21 (3.4) |

